# *Mycobacterium tuberculosis* modifies cell wall carbohydrates during biofilm growth with a concomitant reduction in complement activation

**DOI:** 10.1101/2021.03.23.436651

**Authors:** Thomas Keating, Samuel Lethbridge, Stephen R. Thomas, Luke J. Alderwick, Stephen C. Taylor, Joanna Bacon

## Abstract

There is an urgent need for drugs, new vaccines, and diagnostics for TB. It is recognised that research needed for the development of new vaccines for TB needs to be underpinned by understanding both the molecular and cellular mechanisms of host-pathogen interactions and how the immune response can be modulated to achieve protection with the use of a new vaccine for TB. Complement interacts with and orchestrates many aspects of the innate and adaptive immune responses and activation by *Mycobacterium tuberculosis* can be triggered by all three pathways. However, little is known about the contribution of each of these pathways during TB disease, particularly with respect to mycobacterial phenotype. There is strong evidence for extracellular communities of *M. tuberculosis* during TB disease (biofilms) that are found in the acellular rim of granulomas. These biofilms have been observed in cavities in lung resections from TB patients and are likely to be present in post-primary TB episodes in necrotic lesions. Our study aimed to understand more about the interactions between *M. tuberculosis* biofilms and complement activation, to determine which mycobacterial cell wall components are altered during biofilm growth, and how their alteration contributes to modulation of the complement response. We show that the lectin pathway has a reduced role compared to the classical pathway in initiating complement activation in biofilm bacteria. Analyses of the *M. tuberculosis* biofilm cell wall carbohydrate fractions revealed that there was reduced α-glucan compared to planktonically-grown bacteria. Reduced C3b/iC3b deposition directly onto biofilm carbohydrates was observed which was consistent with both the observed reduction of C3b/iC3b deposition on biofilm bacilli and a reduction in the contribution of the lectin pathway in initiating complement activation on whole bacteria from biofilms, compared to planktonically-grown bacteria.

## Introduction

Tuberculosis (TB) is the number one infectious killer world-wide, responsible for 1.4 million deaths in 2019 and accounts for 25% of all deaths associated with antibiotic-resistance.^1^ There is an urgent need for more effective TB treatments and the disease is challenging to treat, because of host-pathogen interactions and diverse pathologies. If we are to achieve the End-TB goals we need novel, innovative, approaches to understand these interactions and tackle the disease. It has been recognised that the research needed for the development of new vaccines for TB needs to be underpinned by a better understanding of both the molecular and cellular mechanisms of host-pathogen interaction and how the immune response can be modulated to achieve protection with the use of a new vaccine for TB.^2^

Complement interacts with and orchestrates many aspects of the innate and adaptive immune responses and the activation of complement can elicit a variety of immunological effects, which include the stimulation and modulation of T-cell responses (the focus for TB vaccine development) and subsequent control of inflammatory responses.^3^ Complement is pertinent to TB disease as tubercule bacilli will encounter complement proteins in the lung.^4^ Complement activation by *M. tuberculosis* can be triggered by the alternative pathway, classical pathway, and lectin pathway^5^. However, in the lung, the alternative pathway is not significantly activated due to low levels of factor D.^6^ Previous studies have suggested C3b/iC3b deposition onto *M. tuberculosis* is one of the mechanisms utilised by the pathogen to gain access to its intracellular niche.^7^

Complement opsonisation with non-immune serum enhances adherence to monocytes predominantly via complement receptor 3 (CR3).^8^ Furthermore, entry via complement receptors of viable complement-opsonised *M. tuberculosis* enables the pathogen to avoid phagosome-lysosome fusion ^9,10^. Activation of the complement cascade may also provide host protection against *M. tuberculosis* infection. One way is via the release of C5a, which affects the synthesis of key macrophage derived cytokines such as IL-12, IL-1β, TNF-α.^11,12^ While activation of the late complement pathway and subsequent C5b-9 assembly on the pathogen surface may not result in direct lytic activity as observed with some gram-negative bacteria, a study utilising C5 deficient mice demonstrated that C5 is required for bacteriostatic control in *ex-vivo* macrophages and is required for granulomatous inflammation ^3^. Subsequent studies which utilised C5 and C5a receptor-deficient mice also showed that C5 mediated events are essential for generation of a protective granulomatous response against mycobacterial trehalose-dimycolate (TDM).^13,14^ TDM or ‘cord factor’, has an established role in the pathogenesis of *M. tuberculosis* and modulates the maturation of granulomas through C5-mediated events that are critical for responses to TDM.^15,16^ The composition the TDM/TMM and an accumulation of free mycolate have been shown for *in-vitro* pellicle biofilms.^17^ Other cell wall components have also been shown to activate complement. Mannose binding lectin (MBL) and ficolin-3 directly interact with the surface protein Ag85a^18^ and mannosylated lipoarabinomannan.^19^ is known to directly activate the lectin pathway. Despite there being evidence that mycobacterial surface components, such as TDM, can activate complement, the relationship between mycobacterial phenotype and complement activation has not been explored in any depth. *M. tuberculosis* is known to modulate the immune response through its complex lipid and sugar rich envelope.^20–23^ Cell wall modification, due to phenotypic adaptation, such as biofilm growth, is likely to have a role in the alteration of complement activation. TB is a complex, chronic disease, resulting from a plethora of interactions between heterogeneous tubercle bacilli and a variety of host cells in the lung and other compartments of the body. During chronic infection, *M. tuberculosis* persists in biofilms located in extracellular multicellular micro-colonies located in the acellular rim of human and guinea pig granuloma.^24–26^ Our study aims to understand more about the interactions between *M. tuberculosis* biofilms and complement activation and to determine which mycobacterial cell wall components that are altered during biofilm growth, contribute to modulation of the complement response. We used human complement deposition assays to determine which pathways were altered in their activation, in biofilms compared to planktonically-grown *M. tuberculosis*.^27^ There is no significant alternative pathway activity in the lung ^28^. To reflect this, samples were incubated with 10% complement where all pathways are active and with 2% complement where the alternative pathway is inhibited.^6^ Lipids and carbohydrates were differentially extracted, from the two bacterial phenotypes, and biochemical analyses were performed to determine their composition. Specific interactions between extracted cell wall components and different complement activators were explored using quantitative ELISA.

## Results

### There is reduced complement activation by biofilm bacteria compared to planktonic bacteria

The impact of biofilm growth on complement activation was determined in pellicle-grown *M. tuberculosis* by comparing three independent replicate biofilm or planktonic growths. IgG-depleted pooled human plasma was used as a complement source and two-colour flow cytometry analyses used to quantify complement-deposition onto the bacterial surface.^27^ The complement source was used at 10% or 2% (final concentration in the assay) in order to assess activation in either the presence of all three pathways (10% complement) or the presence of only the classical and lectin pathways (2% complement). The absence of the alternative pathway, activation at 2% complement is due to lack of available Factor D.^29^ A reduction in the magnitude of C3b/iC3b deposition (Fig. 1A) was observed on biofilm bacteria compared to planktonic bacteria in both the presence (10% complement P= 0.023) and absence (2% complement P= 0.008) of the alternative pathway. Biofilms also resulted in significantly reduced deposition of C5b-9 (Fig. 1B) at both complement concentrations (10% complement P= 0.003, 2% complement P= 0.0003). However, the difference was greater in the absence of the alternative pathway, which is a physiological condition relevant to the lung.

**Figure 1.**
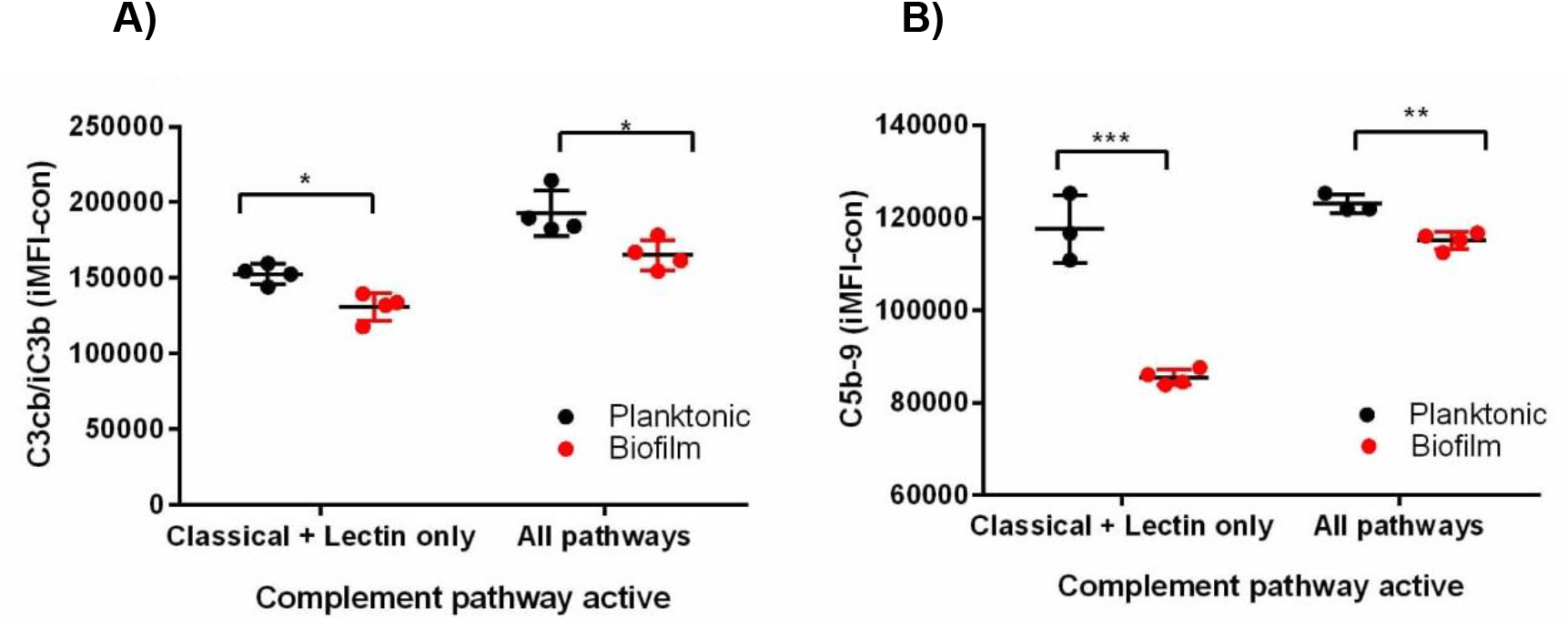
Complement activation by planktonic and biofilm phenotype *M. tuberculosis*. Mean net deposition on planktonic and biofilm *M. tuberculosis* for C3b/iC3b (A) and C5b-9 (B).

### Complement deposition on biofilm grown *M. tuberculosis* is more dependent on the classical pathway than planktonic bacteria and biofilms show reduced lectin pathway activation

The contribution of individual activation pathways to the deposition of C3b/iC3b on the surface of *M. tuberculosis* was assessed. By blocking C1q using a specific monoclonal antibody and therefore preventing classical pathway activation, deposition of C3b/iC3b (Fig. 2) was partially inhibited (P< 0.05). This reduction appeared to be greater in the absence of the alternative pathway (P< 0.02). Despite this reduction, significant C3b/iC3b deposition was observed in the absence the classical pathway (minus C1q) and the alternative pathway on both planktonic and biofilm bacilli show the significance of the lectin pathway for C3b/iC3b deposition on tubercule bacilli. In the absence of the alternative pathway, approximately a third of the C3b/iC3b deposited on planktonic cells is classical pathway-mediated, while the classical pathway contributes two-thirds on biofilm cells; as evidenced by the loss of C3b/iC3b deposition in the absence of C1q. Taken together, these results suggest that the lectin pathway has a reduced role compared to the classical pathway in initiating complement activation onto biofilm cells.

**Figure 2:**
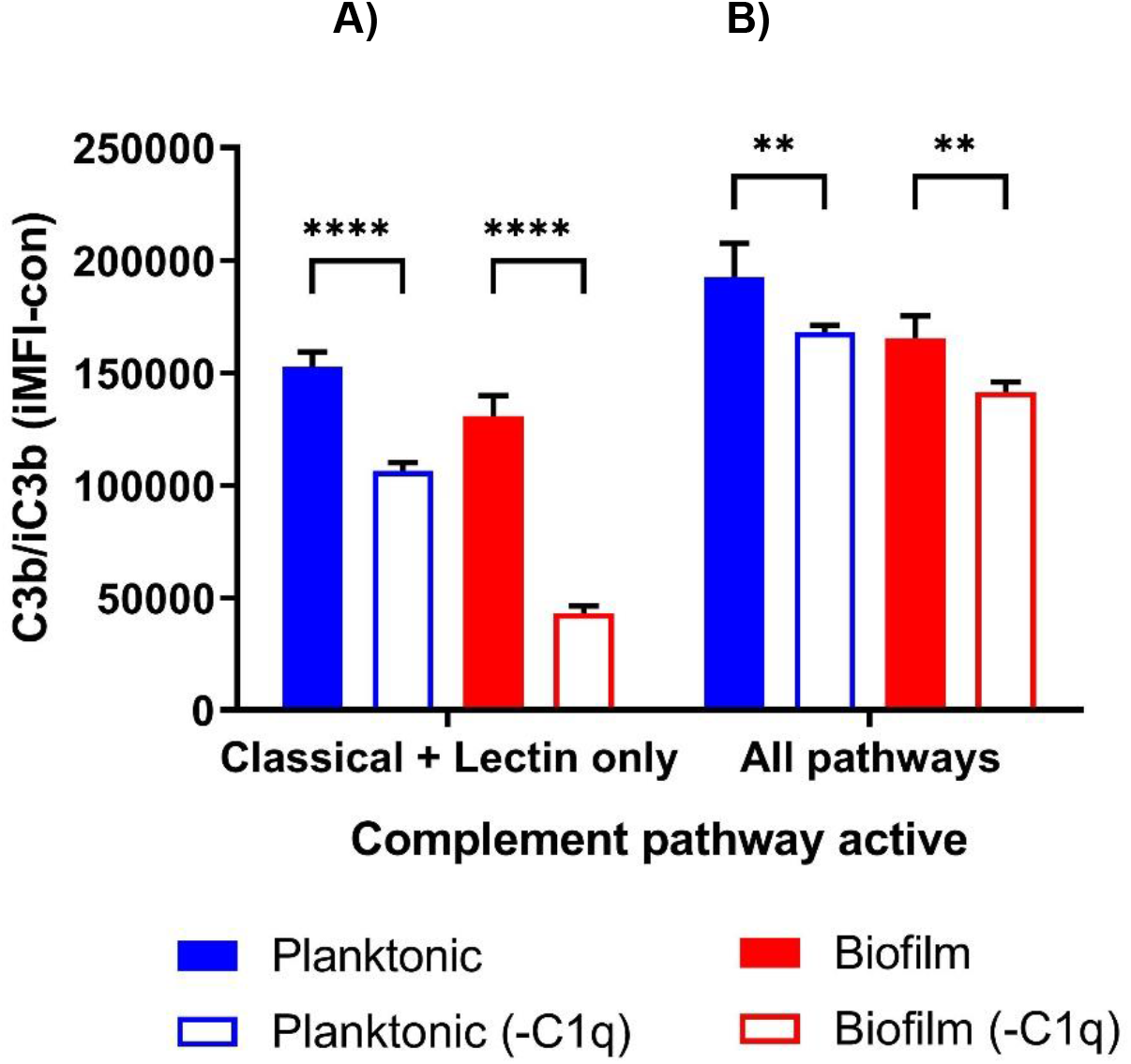
Complement activation by planktonic and biofilm *M. tuberculosis*. Mean net C3b/iC3b deposition without (shaded bars) or with (unfilled bars) pre-incubation with C1q monoclonal antibody (inhibition of the classical pathway) at 2% complement (A) or 10% complement (B). Error bars show standard deviation (n=4)

### There is reduced binding of MBL and C1q to *M. tuberculosis* biofilms

We subsequently quantified the direct binding of classical and lectin pathway activators (C1q and MBL respectively), to planktonic and biofilm *M. tuberculosis* bacteria. In addition, we investigated the binding of Ficolin-3, which is a key ficolin and activator of the lectin pathway and is present in the human lung.^3^ MBL deposition was reduced on biofilm *M. tuberculosis* by 58.9% compared to planktonic bacteria (P = 0.018, Figure 3A). Similarly, direct binding of C1q was also reduced in the biofilm phenotype (P = 0.046 Figure 3B). Ficolin-3 deposition was also reduced on biofilms, but this was not a statistically significant (P = 0.084) (Figure 3C).

**Figure 3.**
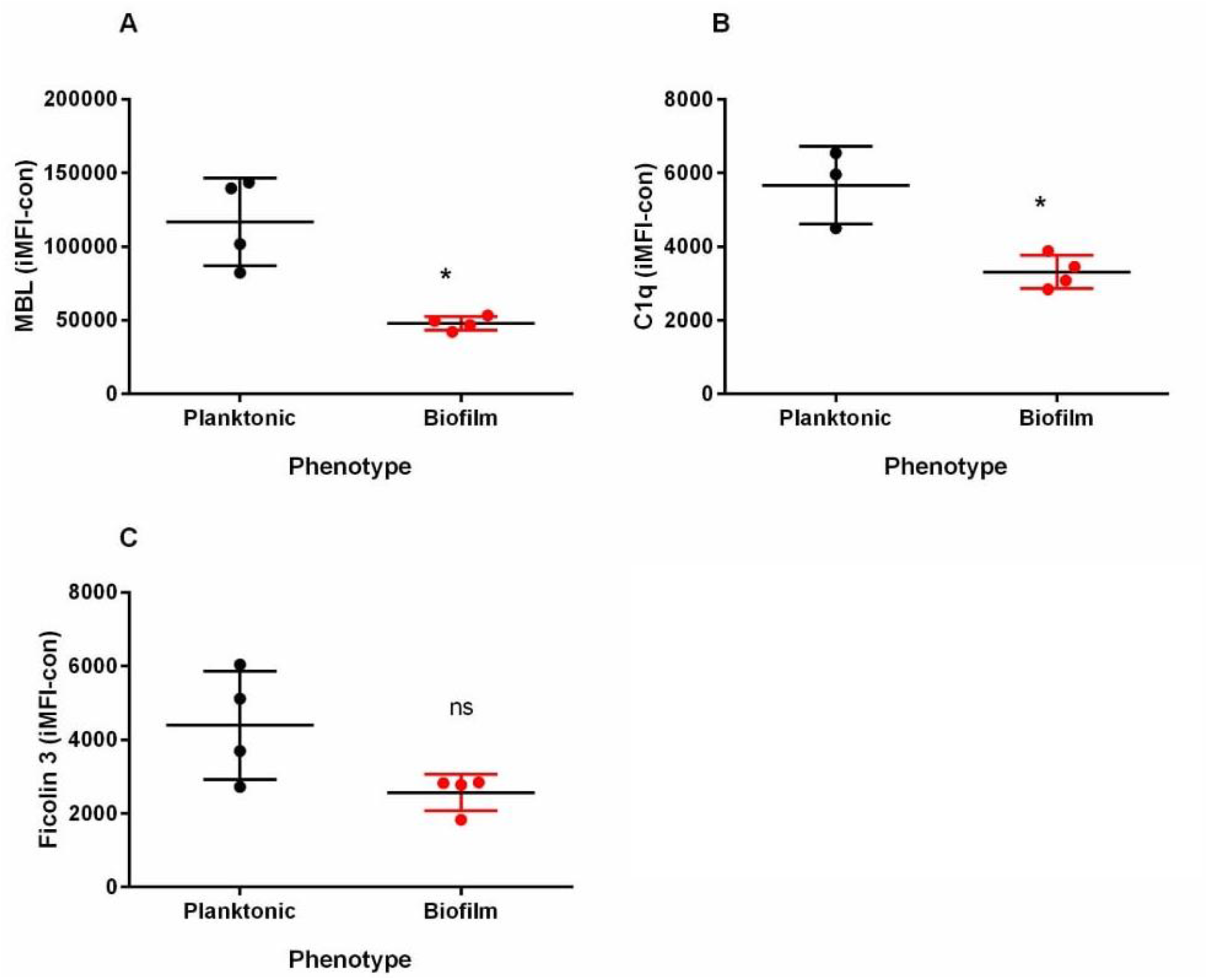
Binding of complement activators to planktonic and biofilm *M. tuberculosis*. Mean net binding to planktonic or biofilm *M. tuberculosis* of mannose lectin binding protein (MBL) (A) C1q (B) or Ficolin-3 (C). Error bars show standard deviation (n=3)

### Biofilm-grown *M. tuberculosis* has an altered carbohydrate composition in the cell wall

We observed the accumulation of extracellular matrix (ECM) in biofilms by scanning electron microscopy (Figure 4.). Analyses of the total lipids confirmed previous findings that there was an accumulation of free mycolic acids. Cell wall-associated carbohydrates were also extracted. The relative proportions of the constituent sugars in the carbohydrate extracts and mycolyl-arabinogalactan-peptidoglycan complex (mAGP) were determined by chemically modifying them to produce alditiol acetate derivatives, which were subsequently analysed by gas chromatography as described previously.^30,31^ The relative proportions of arabinose, mannose, and glucose, from two independent total sugar analyses of planktonic carbohydrate extracts were 17.83±8.65 %, 33.75±0.39 %, and 48.43±8.26 %, respectively. We observed a different pattern in the relative proportions of the sugars for biofilm carbohydrate extracts; arabinose (34.89±0.65 %), mannose (50.33±6.99 %), and glucose (14.78±6.34 %). To gain a clearer understanding of these constituent sugar changes, sugar: sugar ratios were determined (Figure 5A). The proportion of glucose relative to arabinose and mannose was lower in the biofilms compared to planktonic-grown cells, while the proportion of arabinose relative to mannose remained unchanged (Figure 5A). Ratios for rhamnose: arabinose, rhamnose: galactose, and arabinose: galactose in cell wall arabinogalactan were equivalent for the two phenotypes (Figure 5B). These also served as an internal control by confirming that equal quantities of biomass from the two cell states, were used in these comparisons. Taken together, these results show a reduction in glucose in the *M. tuberculosis* biofilm carbohydrate extracts compared to extracts from planktonically grown bacteria. As α -glucan is the only cell wall sugar (captured by these carbohydrate extractions/analyses) to contain glucose, it follows that the prevalence of α-glucan is diminished during the formation of mycobacterial biofilms under the pellicle growth conditions, described in this study.

**Figure 4.**
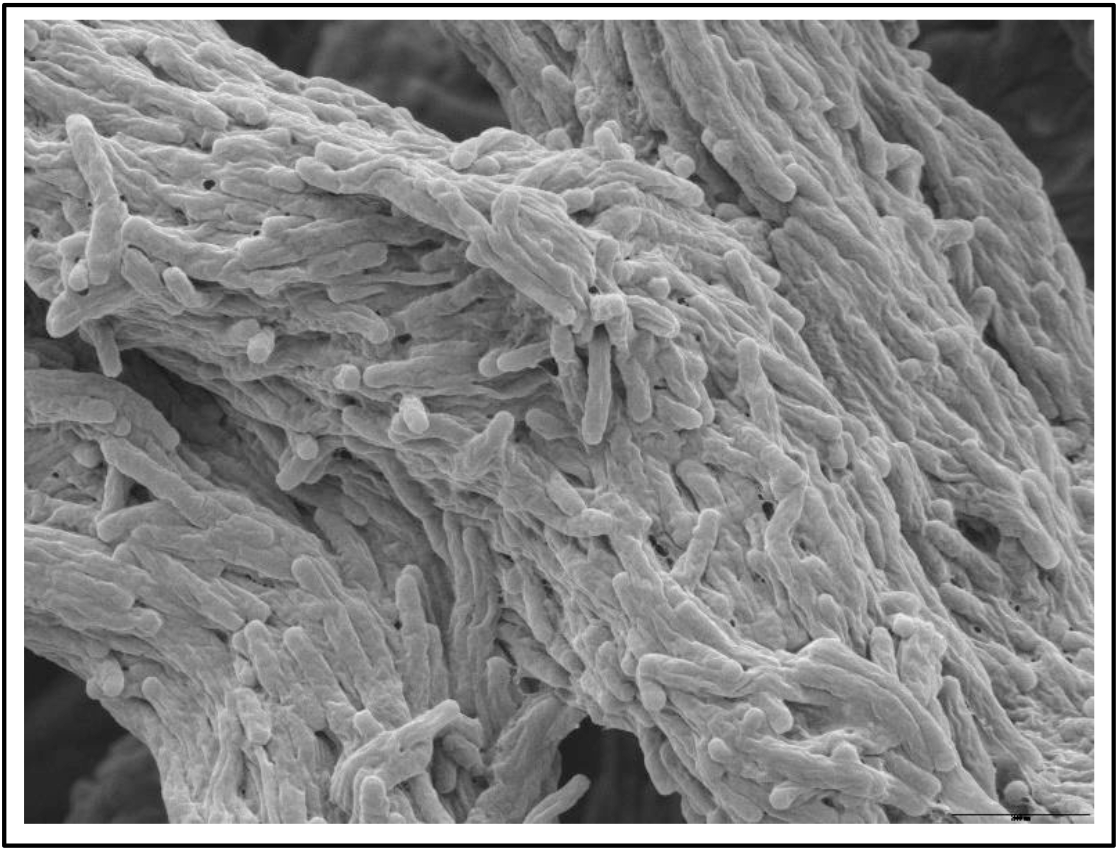
Scanning electron microscopy of M. tuberculosis pellicle biofilms at 5 weeks of growth. Magnification x100

**Figure 5.**
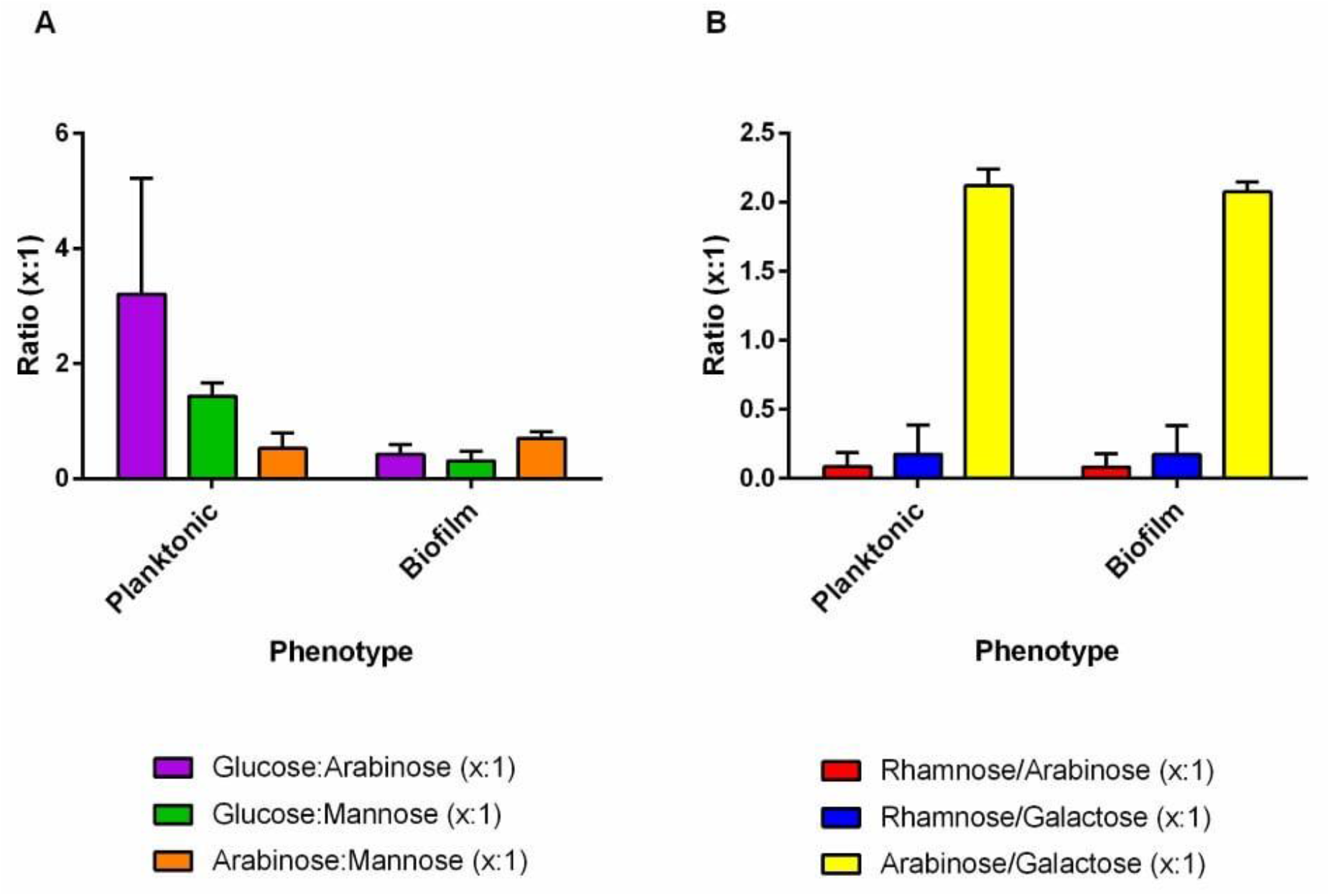
Sugar analysis of M. tuberculosis planktonic and biofilm phenotype carbohydrate and mAGP cell wall extracts. (A) Mean constituent sugar: sugar ratios of planktonic and biofilm phenotype *M. tuberculosis* carbohydrate extracts. (B) Mean constituent sugar: sugar ratios of planktonic and biofilm phenotype mAGP cell wall extracts. Error bars show standard deviation (n=2).

### Deposition of C3b/iC3b was significantly reduced on carbohydrates extracted from biofilm compared to planktonic *M. tuberculosis*

To determine how the different α-glucan levels in biofilm extracts modulate complement and which pathways were differentially activated, we measured the level of C3b/iC3b deposition on carbohydrate extracts from biofilms compared to planktonic growth using a quantitative enzyme-linked immunosorbent assay (ELISA). There was less C3b/iC3b deposition on biofilm carbohydrates compared to planktonic carbohydrates, though this difference was only observed in the absence of the alternative pathway (P= 0.043, Figure 6A) and not when all three pathways were active (P= 0.433, Figure 6C), suggesting that additional amplification provided by the alternative pathway may have saturated the discernible differences in activation between the biofilm and planktonic carbohydrates. The reduced C3b/iC3b deposition elicited by biofilm carbohydrates is commensurate with both the observed reduction of C3b/iC3b deposition on biofilm bacilli and a reduction in the contribution of the lectin pathway in initiating complement activation on whole bacteria from biofilms, compared to planktonically grown bacteria. This highlights the role of carbohydrate alterations during biofilm growth of *M. tuberculosis* and subsequent modulation of the innate immune response.

**Figure 6.**
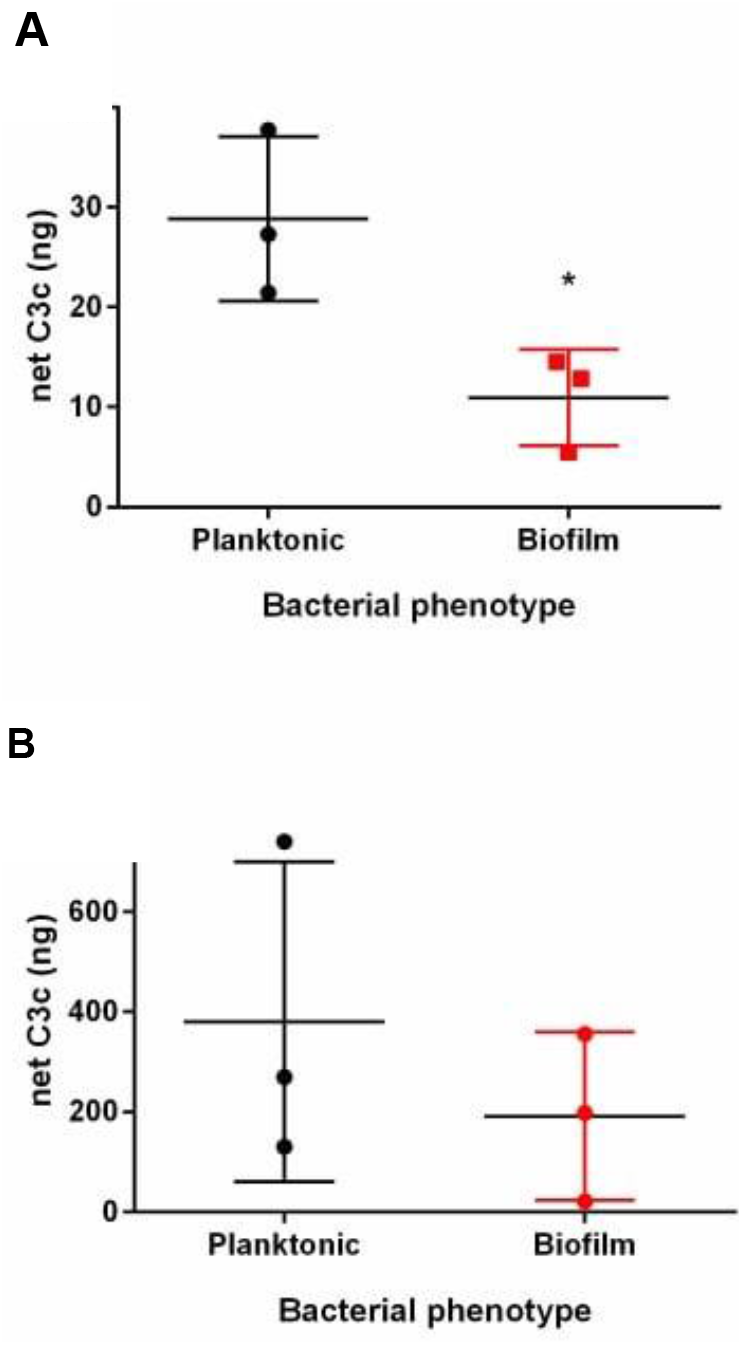
Quantitative ELISA measuring complement deposition on *M. tuberculosis* carbohydrate extracts. C3b/iC3b deposition on planktonic and biofilm carbohydrate extracts incubated with 2% complement (all pathways active) (A) or 10% complement (alternative pathway inactive) (B)

## Discussion

### There is reduced complement deposition on biofilm bacteria (C3b/iC3b and C5b-9)

Previous studies have suggested C3b/iC3b deposition on *M. tuberculosis* is one of the mechanisms utilised by the pathogen to gain access to its intracellular niche^7^ and that entry via complement receptors enables the pathogen to avoid phagosome-lysosome fusion.^8–10,32,33^ In addition to the intracellular phase, there is much evidence now that *M. tuberculosis* persists in an extracellular state in the periphery of cavitary lesions. These extracellular communities are found in the acellular rim of granulomas, have been observed in cavities in lung resections from TB patients, and are likely to be present in post-primary TB episodes in necrotic lesions.^24–26^ Nyka *et al*. demonstrated that these extracellular bacilli resemble biofilms formed by other pathogenic bacteria that cause extracellular infection.^34–36^ These extracellular *M. tuberculosis* communities resemble bacterial biofilms. Extracellular, drug-tolerant, bacilli, have also been observed in drug-treated animals, such as guinea pigs that develop necrotic granulomas.^37–39^ Further confirmation is needed as to whether drug-treatment induces the presence of these biofilm-like communities. The extracellular matrix (ECM) around these acellular rims, typical of biofilms, has been shown to contain *M. tuberculosis* mycolic acids, likely secreted by bacilli.^40^ *M. tuberculosis* can alter these outer-surface molecules and orchestrate a variety of modulated host-responses that dictate the outcome of disease, and the effectiveness of treatment. We need to understand more about this mycobacterial phenotype, other cell wall components that are modified in a biofilm, and their host-pathogen-interaction, so that we can target biofilms with new treatment strategies. With a focus on complement, in this study, the reduction in complement deposition and binding of MBL and C1q on pellicle biofilms, suggest that the organism could be resisting phagocytosis through reduction in the binding of opsonin C3b. Further to this, a reduction in C3b generation and binding would also lead to a reduction in the release of the potent immunomodulatory anaphylatoxins C3a and C5a, in turn, regulating excessive inflammation and activation of cellular responses, which warrant further investigation, as possible targets for the control of necrotic inflammation.

### Complement activation on biofilm *M. tuberculosis* is *more dependent on the classical pathway than for planktonic bacteria and shows reduced lectin pathway activation*

Irrespective of phenotype, we have observed that activation of the complement system by *M. tuberculosis* can occur via antibody-independent activation of classical pathway. The current paradigm is that antibodies are required for classical pathway activation. However, we have shown that for *M. tuberculosis* the classical pathway can be initiated by direct binding of C1q to the cell surface. C1q was blocked using a specific anti-C1q monoclonal antibody and there was a significant reduction in C3b/iC3b deposition on both biofilm and planktonic bacteria, this was especially pronounced on biofilm bacteria. Raised C1q levels have been measured in patients with TB^41^ as compared to other respiratory diseases such as pneumonia and sarcoidosis. Direct binding of C3 either via the classical pathway or the alternative pathway (depending on the sera concentration) has been observed using sera or bronchial lavage from non-immune patients.^42^ Taken together, these data are indicative of a role for the classical pathway in TB disease.

We also observed activation of the lectin pathway by selective inhibition of the alternative and classical pathways and by direct binding of MBL and ficolin-3 to the planktonic or biofilm bacilli. This is supported by previous evidence that activation of the lectin pathway can occur through interaction of cell surface protein Ag85a and lipoarabinomannan with MBL and ficolins.^18,19^With such a carbohydrate-rich cell surface it is unsurprising that *M. tuberculosis* activates the lectin pathway. What has not been shown previously is the impact of growth phenotype on the modulation of lectin pathway activation. There is a significant reduction in MBL-binding on biofilm bacteria compared to planktonic bacilli with a concomitant reduction in lectin pathway activation. Biofilm formation has been associated with reduced activation of the complement system in another respiratory mycobacterial species such as *Mycobacterium abscessus* and there are examples of other bacterial biofilms, that demonstrate modulated complement activation, particularly reduced C3b deposition, via a variety of pathogen-specific mechanisms.^43^ Deposition of C3b and C1q-binding to the bacterial surface of *Streptococcus pneumoniae* biofilms is impaired, enabling pneumococcal biofilms to avoid the activation of the classical pathway. *Pseudomonas aeruginosa* growing planktonically resulted in a stronger activation of complement than biofilms,^44^ which could be inactivating the complement system by secreting alkaline protease and elastase.^45^ Biofilms are likely to have a plethora of wide-reaching interactions and effects on the innate immune response. *Staphylococcus pneumoniae* biofilms also impair phagocytosis and this may be due to the reduction of C3b deposition. We know that complement deposition enables phagocytosis of *M. tuberculosis*. However, we do not yet know the impact of a biofilm phenotype on the interplay between complement, cell-mediated immunity, and the adaptive humoral response.

In our study, and in the patient studies described, it is challenging to determine the contribution of each pathway, as both the classical and lectin pathways are innately activating complement. Following a similar approach to the inhibition of the classical pathway by blocking C1q, we would need to inhibit the lectin pathway in order to understand the specific contribution that each pathway is making. This could potentially be achieved by saturating (thereby by blocking) the activation of MBL complement with mannose. However, this would not completely inactivate the lectin pathway as we have also shown binding of Ficolin 3, which will not bind to mannose and therefore remain active.^46^

### *M. tuberculosis* biofilms show altered complement activation through the modification of α-glucan

To investigate the physiological differences between biofilm and planktonic bacilli that result in altered complement-pathogen interactions, we explored the differences in the composition of cell surface components. We observed changes in the lipids composition and saw an accumulation of free mycolate; the association of free mycolate with the extracellular material of mycobacterial pellicles is now widely accepted.^17^ Polysaccharides are often associated with the extracellular matrix in other bacteria.^47^ However, there have been very few studies that describe carbohydrate changes in the *M. tuberculosis* biofilm. Given the role of bacterial sugars in the activation of the lectin pathway and the differences in the contribution of this pathway to complement activation between the two phenotypes, we chose to focus on the analysis of carbohydrate changes in the cell wall. The outermost non-covalently bound cell wall carbohydrate/lipoglycan fractions were analysed (so could also contain ECM carbohydrates) and highlighted a reduction of a glucose-containing carbohydrate in the biofilms; this is likely to be α-glucan given the extraction process that was used here. Further work is in progress to determine the precise biochemical composition of these carbohydrates. No changes were observed in the proportion of arabinose: mannose confirming that there were no alterations in the lipoglycan composition or arabinomannan between the two phenotypes, such as elongation of the arabinan chain, as observed under a nutrient starved non-replicating state.^30^ Previously, a loosely bound capsule has been associated with *M. tuberculosis*, comprising carbohydrates, particularly α-glucan,^48^ which *is* expressed both *in vitro* and *in vivo*.^49^ These capsular polysaccharides, including α-glucan, will mediate the non-opsonic binding of *M. tuberculosis* H37Rv to CR3,^50^ which may be favourable to the intracellular survival of the tubercle bacillus. However, the role of capsule in pathogenesis has not been determined. Here, the reduced C3b/iC3b deposition elicited by biofilm carbohydrates was consistent with both the observed reduction of C3b/iC3b deposition on biofilm bacilli and a reduction in the contribution of the lectin pathway in initiating complement activation on whole bacteria from biofilms, compared to planktonically grown bacteria. This highlights the role of carbohydrate alterations during biofilm growth of *M. tuberculosis* and the subsequent modulation of the innate immune response. Although it is challenging to directly compare the levels of C3b/iC3b deposition between the experiments with whole bacteria (Fig. 1) and carbohydrates, it appears that the differences in activation between biofilms and planktonic bacteria were greater on whole bacteria. This could be explained by the contribution of other bacterial components (such as cell wall proteins and lipids), to the response present in whole bacteria, and not in the carbohydrate fractions. Given that C3b/iC3b deposition onto *M. tuberculosis* is one of the mechanisms utilised by the pathogen to gain access to its intracellular niche^7^ and that entry via complement receptors enables the pathogen to avoid phagosome-lysosome fusion, reduction of α-glucan may be a mechanism by which *M. tuberculosis* avoids uptake during an extracellular phase of its life cycle. If we also consider these extracellular bacterial biofilms in the context of TB transmission; expectorated *M. tuberculosis* biofilm bacteria could be entering the lung environment of a new host where the alternative pathway is not active.^51^ Complement activation leads to the release of the anaphylatoxin C5a, which is required for tuberculosis control.^51^ Reduced of complement activation by these bacteria may provide an advantage in the establishment of infection.

### Concluding remarks

Our study reveals that *M. tuberculosis* cultured as a pellicle biofilm leads to reduced complement activation. We show here that *M. tuberculosis* biofilms modify their cell wall carbohydrates. Consistent with this finding is reduced classical and lectin pathway activation, which is associated with reduction of C3b/iC3b deposition on biofilm carbohydrate extracts We have highlighted the role of carbohydrate alterations during biofilm growth of *M. tuberculosis* and subsequent modulation of the innate immune response, which may be one mechanism by which, *M. tuberculosis* avoids phagocytosis in establishing an extracellular state in the lung.

## Experimental Procedures

### *M. tuberculosis* planktonic and biofilm culture

All cultures were grown in modified Sauton’s medium with the following ingredients: KH_2_PO_4_ 0.05 mg mL^-1^, MgSO_4_ 7H_2_O 0.5 mg mL^-1^, Citric acid 2.0 mg mL^-1^, Ferric ammonium citrate 0.05 mg mL^-1^, Glycerol 75.6 mg mL^-1^, Asparagine 4.0 mg mL^-1^ and adjusted to pH 7.4). The inoculum used for all cultures (planktonic and biofilms) came from steady-state fast-growth chemostat cultures grown in CMM MOD2 medium.^52,53^ Planktonic cultures were grown in aerated flasks shaken at 200 rpm at 37°C for 7 days, from a starting optical density of 0.05 OD_540nm_. Biofilms were grown in 24-well plates. These were inoculated using starter cultures, containing modified Sauton’s medium that had been inoculated with chemostat culture to an optical density of 0.05 OD_540nm_ and grown for 7 days shaken at 200 rpm at 37°C. Each well was inoculated with 2mL of culture. These were incubated for 5 weeks, statically, at 37°C in an airtight container. Planktonic cultures were harvested by pelleting at 3060g for 10 minutes and biofilm cultures were harvested by scraping biomass from the air-liquid interface using sterile disposable spatulas, into a glass tube. Cell biomass was dried down, using a Genevac EZ-2 plus evaporator, in preparation for extractions. An equivalent quantity of biomass from each culture type was used for further extractions and complement deposition assays. Biomass was quantified by viable count and/or dry weight, depending on the analyses. Generally, biomass scraped from the wells of eight plates were pooled to give a single replicate biofilm sample.

### Dispersion of *M. tuberculosis* cultures

To provide biomass samples for complement deposition and analyses by flow cytometry, biomass was dispersed by agitation with 4 mm glass beads. Briefly, live *M. tuberculosis* planktonic pellets or scraped biofilms were agitated with sterile 4 mm glass beads which had been added at a ratio of approximately 5:1 bead volume: pellet volume. The samples were mixed using a vortex for 1 minute, suspended in PBS and left to sediment for 10 minutes. The liquid was poured into fresh tubes and the suspension was spun at 200 g. The supernatant containing the dispersed cells were aliquoted into cryovials and frozen at −80°C until required.

### Scanning Electron Microscopy of intact *M. tuberculosis* biofilms

*M. tuberculosis* biofilms were cultured in 24-well plates (as described above) containing sterile plastic coverslips, cut to size, and placed into each well. After 5 weeks of biofilm growth the coverslips were removed and fixed in 4% formaldehyde (v/v in water). Ethanol/solvent dehydration: Formaldehyde was carefully removed and replaced with 2% osmium tetroxide for 2 hours at room temperature for secondary fixation. Biofilms were then dehydrated through graded ethanol solutions for 15 minutes at room temperature sequentially at 25%, 50%, 75%, 100% and 100% concentration. Following this, the coverslips were dehydrated with 100% Hexamethyldisilane for 15 minutes at room temperature; this step was repeated. Coverslips were then air-dried and mounted on SEM stubs using double-sided adhesive carbon discs. The mounted coverslips/pellicles were then conductive coated with approximately 10nm thickness of gold using an ion beam sputter coater (AtomTech 700 series Ultra Fine Grain Coating Unit).

### Non-covalent carbohydrate and mAGP extraction

The method described by Besra *et al*., was followed.^54^ Briefly, biomass was harvested from three replicate planktonic or biofilm *M. tuberculosis* cultures, inactivated by autoclaving at 126°C for 30 minutes and evaporated to dryness using a Genevac EZ-2 plus evaporator. Planktonic or biofilm biomass was heated under reflux with 10 mL ethanol-water (1:1) at 75°C for 4 hours. Samples were then left to cool to room temperature, spun at 3000g for 15 minutes and the supernatant was removed into fresh tubes. The pellet was topped up to with 10 mL ethanol-water (1:1) and the heating step and centrifugation was repeated. The supernatants were vacuum-dried, and the pellets were recombined in phenol saturated with PBS. The samples were heated at 75°C for 30 minutes and left to cool to room temperature. The phenol and aqueous layers were separated by centrifugation at 3000 g. The aqueous layer was removed into a semi-permeable dialysis membrane (MWCO 3500). The samples were dialysed overnight with running tap water and left in distilled water for 1 hour to remove salts from the tap water. Samples were transferred into clean pre-weighed glass tubes and were subsequently dried. The remaining pellet from the ethanol reflux was used for mAGP extraction. 2% SDS (w/v in PBS) was added to planktonic and biofilm pellets that were remaining from the ethanol reflux step of the carbohydrate and lipoglycan extraction. Samples were heated under reflux at 95°C overnight. Following this, the samples were washed with water, pelleted, washed with 80% acetone (v/v in water), pelleted, washed with 100% acetone, and dried.

### Total sugar analysis

Method as described in ^55,56^. Planktonic and biofilm phenotype carbohydrate and mAGP extracts were treated with 2M Trifluoroacetic acid (TFA). 200 μL of 2M TFA was added to 0.5 mg mAGP or carbohydrate extracts. Samples were heated to 120°C for 1.5 hours under reflux and allowed to cool to room temperature. The acid was evaporated using a sample concentrator. 100 μL of 10 mg mL^-1^ NaBH_4_ in 1:1 ethanol: NH_4_OH (1M) was added to each sample and left capped at room temperature overnight. 3 drops of glacial acetic acid were added to each sample and evaporated. Three drops of 10% glacial acetic acid in methanol were added then evaporated. This step was repeated once. Three drops of 100% methanol were added and evaporated. This step was repeated. 100 μL acetic anhydride was added and this was heated at 120°C for 1 hour. After allowing the sample to cool to room temperature, 100 μL of toluene was added to the samples and this was evaporated. 2 mL of chloroform and 2 mL of water were added to each sample. The lower organic layer was transferred into a fresh tube using a glass pipette and dried for gas chromatography analysis. The percentage of constituent sugars was determined by calculating the relative area under arabinose, mannose, glucose peaks for carbohydrate/lipoglycan fractions and by calculating the relative area under rhamnose, arabinose and galactose peaks for mAGP extract from planktonic and biofilm cultures. Sugar: sugar ratios were calculated from the total sugar analysis.

### Quantitative ELISA

A previously published ELISA method was used with modifications.^57^ Three biological replicates of planktonic or biofilm *M. tuberculosis* carbohydrate extracts at 15 mg mL^-1^ were diluted in 1 mL of carbonate buffer (Sigma carbonate/bicarbonate capsule in 100 mL distilled water) to a final concentration of 25 µg mL^-1^. 100 µL was added to eight wells of a microtitre plate to ensure 2.5 µg of carbohydrate per well. Seven-point standard curves were generated using 1:4 serial dilutions of human purified C3 (CompTech). Microtitre plates were sealed and left at 4°C overnight to bind carbohydrates and protein standards to the bottom of the wells. The buffer was removed by inverting the plate. The plate was blocked by adding 100 µL of 10% BSA (v/v in PBS) to each well and incubating for 1 hour. The wash step was repeated and 100 µL of PBS containing 0.05% Tween 20 was added to each well and washed again. 100 µL of complement binding buffer (CBB; 1.76 mM MgCl2, 0.25 mM CaCl2, 145.4mM NaCl), 2% complement in CBB, and 10% complement in CBB was added to respective wells and incubated for 1 hour at 37°C. Plates were washed as previously described. For C3b/iC3b ELISAs, 100 µL of (1:500) pAb anti-C3c HRP (Abcam) in CBB was added to each well. The plate was sealed and incubated for 1 hour at room temperature. Plates were quantified using a Multiskan EX plate reader at 450 nm with Ascent software. 5 parameter standard curves to quantify bound C3b/iC3b with absorbance values were generated using GraphPad Prism software. Significant differences in C3b/iC3b deposition on planktonic and biofilm phenotype carbohydrate extracts was determined using Welch’s t-test in GraphPad Prism software.

### Dispersion of *M. tuberculosis* cultures

Agitation with 4 mm glass beads was performed as described previously ^55,56^ with the following modifications. Briefly, live *M. tuberculosis* planktonic pellets or scraped biofilms were agitated with sterile 4 mm glass beads which had been added at a ratio of approximately 5:1 bead volume: pellet volume. The samples were mixed using a vortex for 1 minute, suspended in PBS and left to sediment for 10 minutes. The liquid was poured into fresh tubes and the suspension was spun at 200 g. The supernatant (dispersed whole cell fraction) was aliquoted into cryovials and frozen at −80°C until required.

### C3b/iC3b deposition ±mAb C1q assay

Viable, thawed planktonic and biofilm phenotype *M. tuberculosis* from glass-bead treated stocks, were diluted to 0.2 OD540nm in complement binding buffer (CBB). This buffer was made by dissolving a complement fixation diluent tablet (Oxoid, UK) in PBS containing 2% bovine serum albumen. CBB contained 1.76 mM MgCl2, 0.25 mM CaCl2, 145.4mM NaCl. IgG-depleted exogenous pooled human complement was pre-incubated for 20 minutes at 4°C with PBS (control) or mouse anti-human C1q mAb (Hycult Biotech, Netherlands) at 1/5 dilution. To bacteria/zymosan control wells, 55 μL of CBB was added. To 2% complement wells, 2.5 μL of IgG-depleted exogenous pooled human complement-PBS/mAb anti C1q (4:1) was added along with 52.5 μL of CBB. To 10% complement wells, 12.5 μL of IgG-depleted exogenous pooled human complement-PBS/mAb anti C1q (4:1) was added along with 42.5 μL CBB. The plate was incubated at 37°C for 45 minutes 900 rpm. The plate was centrifuged at 3060g for 5 minutes and each well was washed with 200 μL CBB. The plate was spun again at 3060g and re-suspended in 4% PBS-formaldehyde, to inactive the bacteria. The plate was spun, by centrifugation, at 3060 g and re-suspended in CBB. This spin was repeated and re-suspended in 200 μL of CBB containing FITC rabbit anti-human C3c pAb (Abcam, UK) a 500-fold dilution. The plate was incubated at 4°C for 20minutes, spun at 3060 g for 5 minutes, and washed twice with PBS before being analysed by flow cytometry using a CyAN ADP Analyser flow cytometer (Beckman-Coulter, USA).

After running the plate, compensation was performed using Summit 4.3 software (Beckman-Coulter, USA) from single conjugate sample FCS files and the magnitude of C3b/iC3b deposition was measured by calculating integrated median fluorescence intensity (Darrah et al., 2007) (median fluorescence*% positive cells) for each sample. The iMFI of bacteria + conjugate only negative control wells were subtracted from each sample to remove the contribution of background fluorescence to net iMFI values. For statistical analysis, ≥3 biological replicate planktonic and biofilm samples were compared by performing t-tests corrected for multiple comparisons by false discovery rate ^57^ with Q = 5% using GraphPad Prism software.

### MBL, C1q and Ficolin-3 binding assays

Viable, thawed planktonic and biofilm phenotype *M. tuberculosis* from glass-bead treated stocks, were diluted to 0.2 OD_540 nm_ in CBB. 55 µL of CBB was added to control wells and 54 µL added to MBL/C1q/Ficolin-3 wells. 45 µL of bacteria was added to designated wells. In separate experiments, recombinant MBL (R&D systems) (100 µg mL^-1^) or recombinant ficolin-3 (R&D systems) or human purified C1q (Comp Tech) was added to designated wells to give a final concentration of 1µg mL^-1^ in 100 µL. Plates were incubated at 900 rpm 37°C for 45 min. Plates were spun at 3060g for 5 minutes and washed with 200 µL CBB per well. Plates were spun at 3060g for 5 minutes and re-suspended in either 1:100 mouse anti-human MBL (Hycult), 1:500 mouse anti-human C1q (Quidel) or 1:100 mouse anti-human Ficolin-3 (Hycult) in CBB or CBB only and incubated for 20 minutes at room temperature. Plates were spun at 3060g 5 minutes and washed with 200 µL CBB per well. Plates were spun at 3060g 5min and re-suspended in 1:500 Goat Anti-Mouse IgG (H+L) Fluorescein (FITC)-AffiniPure F(ab’)2 Fragment antibody in CBB or CBB only and incubated for 20 minutes at room temperature. Plates were spun at 3060g for 5 minutes and washed with 200 µL filter sterilised PBS per well and this step was repeated. Plates were spun at 3060g for 5 minutes and re-suspended in 200 µL 4% formaldehyde in PBS per well and sealed with 70% ethanol-soaked plate sealers. Plates were sprayed with 70% ethanol and incubated at room temperature for 1 hour to inactivate the bacteria in the wells. For statistical analyses of the MBL, C1q and ficolin-3 experiments, ≥3 biological replicate *M. tuberculosis* planktonic and biofilm samples were compared by performing two-tailed Welch’s t-tests using GraphPad Prism software.

## Author contributions

Conceptualisation: S.T, J.B. Methodology: S.L, T.K, S.C.T, S.R.T, J.B, and L.J.A. Formal Analysis: T.K, S.C.T. Investigation: S.L, T.K. Resources: S.C.T, J.B, and L.J.A. Writing Original Draft: J.B. Review & Editing, S.T, J.B, T.K, S.L, and L.J.A. Supervision, S.C.T, J.B, and L.J.A. Funding Acquisition, S.C.T, J. B.

## Acknowledgements

This work was funded by Public Health England. The authors declare no conflicts of interest in the publication of this work. We thank Dr Andrew Gorringe for reviewing and helping with the preparation of this manuscript.

## Notes

### Competing Interest Statement

The authors have declared no competing interest.

